# Decoding Olfactory Bulb Output: A Behavioral Assessment of Rate, Synchrony, and Respiratory Phase Coding

**DOI:** 10.1101/2024.10.26.620381

**Authors:** Xiaochen Fu, Janine K. Reinert, Sander Lindeman, Izumi Fukunaga

**Affiliations:** Sensory and Behavioural Neuroscience Unit, Okinawa Institute of Science and Technology Graduate University

## Abstract

The olfactory system is a well-known model for studying the temporal encoding of sensory stimuli due to its rhythmic stimulus delivery through respiration. The sniff rhythm is considered critical to structuring the output of computation in the primary olfactory area, the olfactory bulb. Here, Neuropixels recording from awake, head-fixed mice confirmed that both the rate and sniff-locked spike timing are informative about odour identity. We tested the behavioural importance of these temporal features using simple closed-loop optogenetics embedded in custom behavioural paradigms. We found that mice perceive differences in evoked spike counts and discriminate between synchronous vs. asynchronous activations of the output neurons. Surprisingly, they failed to distinguish the timing of evoked activity relative to the sniff cycle. These results challenge the utility of internally generated rhythms as reference signals in the neural encoding of the environment.

## Introduction

As animals explore their environment, the brain’s sensory systems respond continually to incoming signals with intricate spatial and temporal activity patterns. Studying how the brain encodes external information is important because it reveals how it may optimise its limited resources and how physiological functions emerge from intrinsic properties and synaptic interactions of neurons. The sensory system may optimise the stimulus encoding by matching the neuronal responses to the statistics of the sensory environment, leading to efficient uses of energy or the dynamic range available ^1–6^. This optimisation may also serve to guide animals’ behavioural choices, ultimately to maximise their fitness ^7,8^. A critical step in understanding the mechanics behind such optimization processes is understanding the format the sensory systems use to encode the environment.

Olfaction is a chemical sense that starts with the recognition of odorous molecules by olfactory receptors in the nasal epithelium ^9^. The choice of olfactory receptor, and therefore, the identities of neurons and the subsequent connectivity patterns, are some of the important determinants of the information conveyed ^10–15^.

The olfactory system has also attracted much attention as an ideal system to study the role of timing in encoding sensory stimuli ^16–22^. It is because of widespread sniff-locking in this system, which originates in the fact that olfactory stimuli are airborne, and their delivery couples to the animal’s rhythmic intake of air (sniffs) and its active modulation. This sniff-locking sets the fundamental frequency of activity for most neurons in the early and intermediate stages of olfactory processing. ^20,23–34^. It is particularly the case for the primary olfactory area, the olfactory bulb^16,20,23,28,30,32,35^, which receives the olfactory nerve input and whose outputs are conveyed by at least two classes of principal neurons, mitral cells and tufted cells. The prominent sniff-locking by these olfactory bulb neurons has inspired theories on how the brain may use such timing to encode stimuli. These include the importance of the earliest activity in encoding the identity of odours ^21,36,37^ or encoding olfactory stimuli with latencies in a population of neurons ^19,20,38^, which may have advantages over spike counts (the rate code), for example, in simplifying some computations.

Patterns of activity evoked in the olfactory bulb contain information via both firing rate changes and action potential timing ^23,39^. However, it has been argued that correlating neural features and sensory information on its own does not guarantee the importance of those features in driving behaviour ^7^. A link to behavioural output is needed, ideally through intervention methods by manipulating the activity of neurons experimentally in behavioural experiments. This approach has demonstrated that sniff timing is salient for the periphery: when the olfactory nerve is stimulated optogenetically, mice are sensitive to when, within the sniff cycle, the stimulation occurs ^21,22^. Subsequent optogenetic experiments on the olfactory bulb output indicate that some aspects of timing are important ^40,41^, but it remains an open question if structuring the activity with respect to the sniff rhythm and the firing rate are perceptible features of the olfactory bulb output.

In this study, we hypothesise that both firing rate modulation and precise timing of second-order neurons of the olfactory system are important for perception and driving behaviour. We used optogenetics to stimulate the olfactory bulb output neurons. By precisely triggering activity on respiratory cycles and testing which patterns are discriminable by animals behaviourally, we found that the number of action potentials and relative latencies are accessible but not the timing with respect to the sniff phase.

## Results

### Large-scale electrophysiology demonstrates that the rate and latencies both contain olfactory information

Sniff rhythms set a fundamental temporal structure for the olfactory system. To study how the temporal patterns of the olfactory bulb output relate to sensory information, we presented odours to awake, head-fixed mice that have been habituated but otherwise naïve. We recorded the spiking activity of neurons extracellularly using a high-density neural probe (Neuropixels 1.0 ^42^). The mice received rewards (water droplets) on randomly selected trials, which kept the mice engaged. Odours evoked complex patterns of responses in the recorded olfactory bulb units, as shown in the peri-stimulus time histogram (Fig. 1; Supplemental Fig. 1). Most of the units significantly locked their baseline activity to the sniff rhythm (Supplemental Fig. 2; 274/319 units), indicating a dominant effect this temporal structure has on the activity pattern of olfactory bulb neurons. Indeed, the odour-evoked responses appeared more robustly when the spikes were aligned to the sniff timing rather than the onset of the odour valve opening (Fig. 1D). These observations suggested that both the firing rate and timing may be used to encode olfactory stimuli as previously shown ^23,39^. In our hand, too, a classifier (multi-class support vector machine) trained on both the firing rate or spike latencies decoded the odour identity of test trials well above chance (Fig. 1E; Mean accuracy with a 300 ms window = 53.7 ± 20.0 and 43.8 ± 22.5 for the rate and latencies, respectively; mean ± SD; n = 5 mice). We, therefore, confirm that information about the odour identity is contained in both the rate and timing in the primary olfactory area.

**Figure 1:**
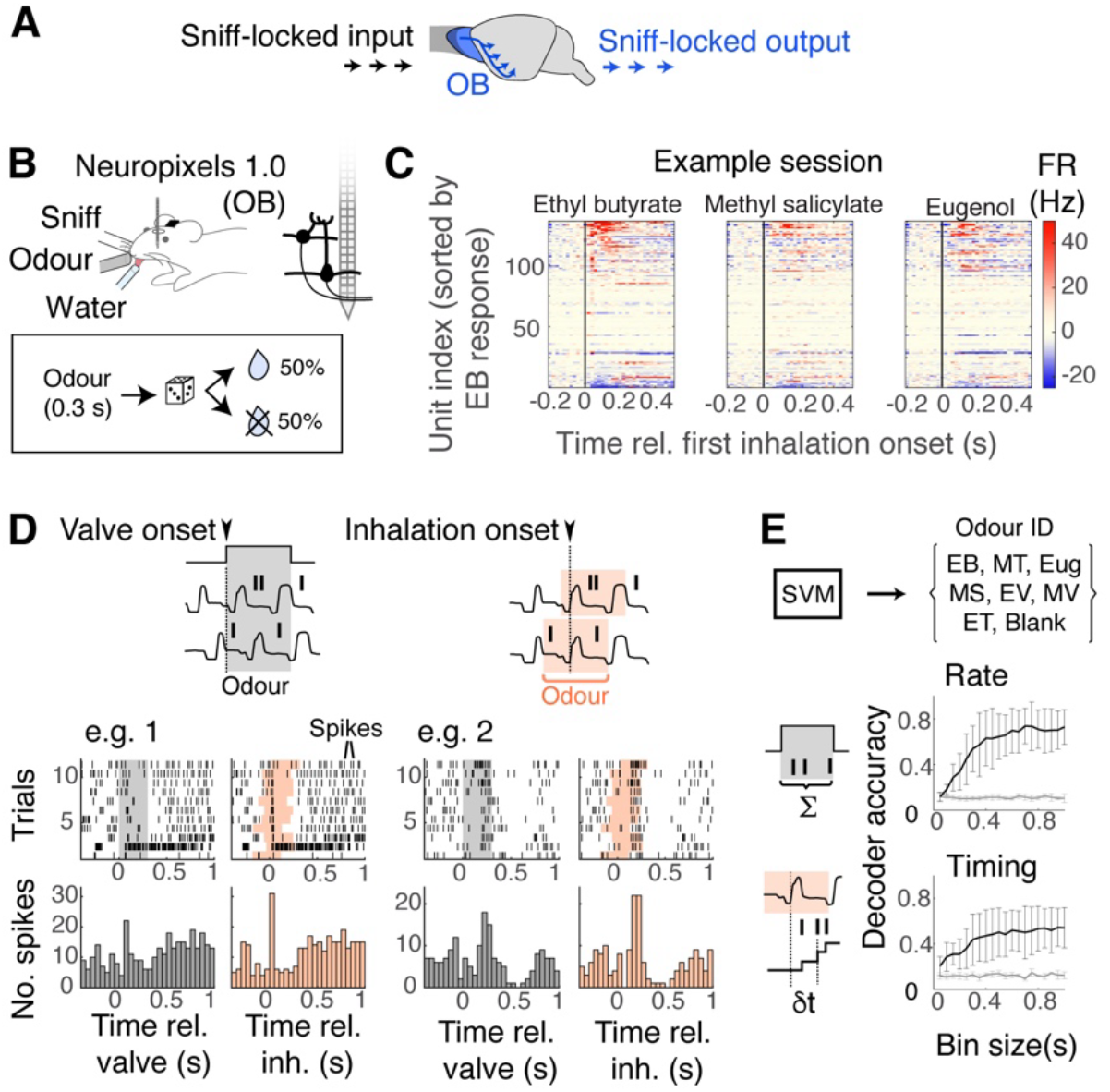
Odour identity information is present in the rate and timing of the olfactory bulb output. **A**. Rhythmic activity in the olfactory bulb (OB) originates in an animal’s sniffing. Both the input and output of the olfactory bulb are sniff-locked. **B**. Experimental configuration. Neuropixels 1.0 probe was inserted into the left olfactory bulb of an awake, head-fixed mouse. 300 ms-long odour presentations were followed by water reward on 50% of the trials to keep the mice engaged. **C**. Peri-stimulus time histograms for simultaneously recorded units from an example session. Colour corresponds to the firing rate change relative to the baseline. **D**. Spike times for each unit were analysed with respect to the valve opening (left) or to the onset of the first inhalation after the valve opening (right). **D** shows spike rasters and peri-stimulus time histograms using the two temporal alignment methods for two example units. **E**. Decoding analysis using a multi-border support vector machine (SVM), using the number of action potentials recorded in the time window (top) or the latency of spikes relative to the inhalation onset (bottom). The accuracy is for test trials (10% of total trials). Trial-shuffled control is shown with grey lines. Odours used = ethyl butyrate (EB), methyl tiglate (MT), Eugenol (Eug), methyl salicylate (MS), ethyl valerate (EV), methyl valerate (MV), ethyl tiglate (ET), and blank air (Blank). N = 5 mice.

**Figure 2:**
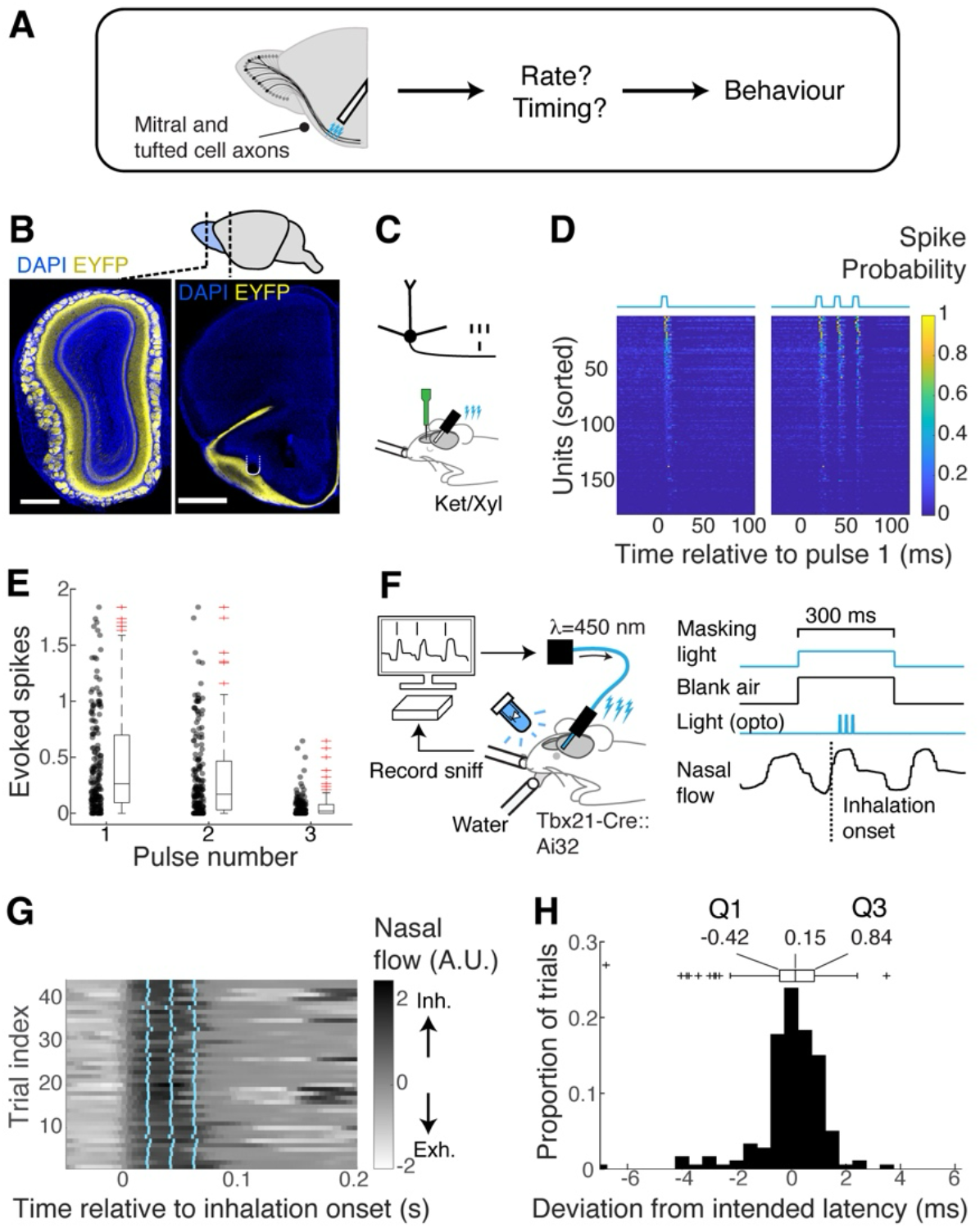
Reliable recruitment of OB output neurons with optogenetics and its closed-loop application. **A**. Schematic showing optogenetic stimulations of mitral and tufted cell axons in the lateral olfactory tract. **B**. Confocal images showing the locations of EFYP-ChR2 expressions in the olfactory bulb (left) and lateral olfactory tract where the optical fibre tip was located (right). Scale bars = 1mm for both. **C**. Schematic. Experimental configuration to verify the optogenetic activation of mitral tufted cells using a 16-channel probe from the olfactory bulb of anaesthetised mice. **D**. Spike probability for each unit from repeated presentations of a single light pulse (left) and 3 pulses of light at 50 Hz (right; 19.2 mW). The pulse duration was 5 ms. **E**. Summary quantification showing the number of spikes evoked in each unit per light pulse. N = 179 units, 4 mice. **F**. Experimental configuration for the closed loop system. Ongoing sniff patterns are measured with a flow sensor adjacent to the nostril, and inhalation onsets are detected online. To mimic an olfactory presentation, the final valve is opened to present blank air, and the first inhalation after the valve opening triggers the light presentation with a specific delay. **G**. Timing of light pulses presented (cyan) shown against the nasal flow measurement for an example session. **H**. Method for calculating the temporal precision (showing one animal). Quartile deviation =(Q3-Q1)/2.

### A simple closed-loop system for activating the olfactory output with precise, within-sniff timing

We designed an experimental system that allows the temporal features of mitral and tufted cell activity to be manipulated in the context of behaviour. We implemented synthetic perception using optogenetics, using a simple closed-loop system, where precise activation of neurons occurs synchronised with the animal’s sniffing, which was analysed online. For the optogenetic stimulation, we used Tbx21-Cre::Ai32 mice, where ChR2-EYFP is expressed in mitral and tufted cells of the olfactory bulb (Fig. 2A,B). Since the olfactory bulb output passes through the lateral olfactory tract, we targeted the stimulation to this axon bundle. This foregoes strong, sniff-locked excitation and inhibition from the local network within the olfactory bulb ^20,28,43^. Our stimulation recruited 31.3% (56/179 units) of recorded olfactory bulb neurons of anaesthetized mice (Fig. 2C-E), and the probability of evoking a spike did not change with sniff latencies (Supplemental Fig. 3; p = 0.996 and 0.953 for putative tufted cells and mitral cells, respectively; F = 0.02 and 0.11; Effect of latency in a 2-way ANOVA test; n = 89 putative TCs and 113 putative MCs, 3 mice). At 50 Hz stimulation, action potentials can be evoked repetitively so that changing the number of light pulses changed the number of evoked action potentials (mean number of evoked action potentials per unit = 0.25 ± 0.29 and 0.79 ± 0.81 for 1 pulse and 3 pulses, respectively; p = 5.45 × 10^−15^, 2-tailed Wilcoxon rank sum test, n = 179 units, 4 mice). There was, however, a slight decline in the efficacy for later pulses (Fig 2E; slope of linear fit with 95% confidence interval = −0.187 ± 0.339, n = 179 units, 4 mice). We also confirmed that our closed-loop system is temporally precise with respect to the sniff cycles in awake mice (Fig. 2H; deviation from the intended timing=0.88 ± 0.38 ms; mean ± SD quartile deviations; n = 12 mice). Overall, these results indicate that we have a reliable method of manipulating the output of the olfactory bulb to test explore the importance of temporal patterns in the context of behaviour.

**Figure 3:**
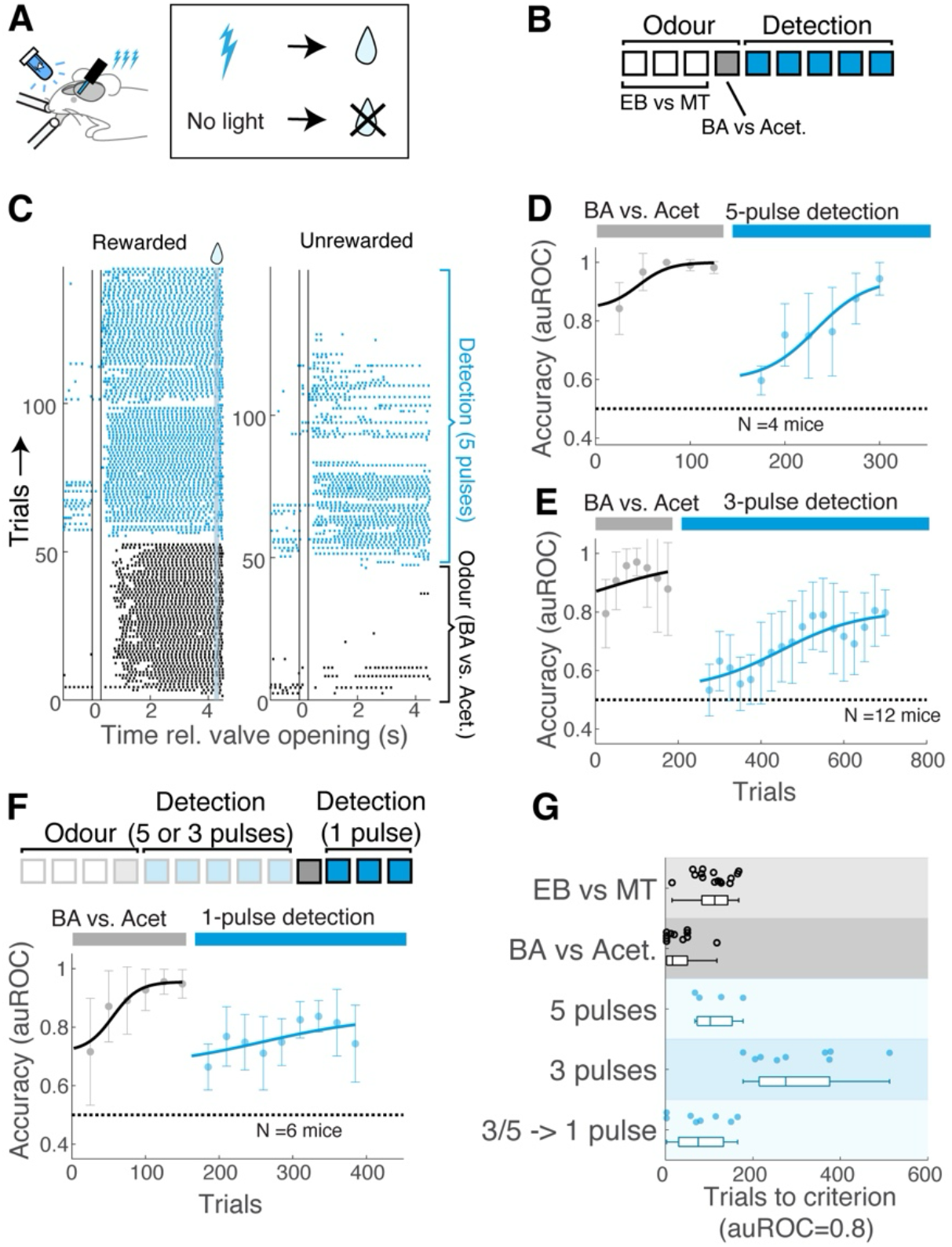
Mice efficiently learn to detect optogenetic activations of OB output. **A**. Experimental configuration for closed-loop optogenetics with Go/NoGo paradigm. In light detection task, water reward was delivered on trials with optogenetic LOT stimulation. **B**. Experimental timeline. The head-fixed mice underwent the light detection task after successfully learning to discriminate two pairs of odours. **C**. Lick raster plots for an example animal. Tick marks indicate the licks times relative to the valve opening and closing (t = 0 and 0.3 s; vertical lines) for rewarded trials (left panel) and unrewarded trials (right panel). Water reward was delivered at 4.3 s after the valve opening. **D**. Learning curve to detect 5 stimulations of LOT (50 Hz) vs. none. N = 4 mice. **E**. As in D but for 3 stimulations. N = 12 mice. **F**. Some mice that learned to detect 3 or 5 stimulations were tested to detect 1 stimulation vs. none (n = 4 mice and 4 mice, respectively). Mean and SD shown. **G**. Summary of learning speed for the odour discrimination (black) and light detection (blue). Vertical lines in the box = 25th, 50th, and 75th percentiles and horizontal whiskers = data range.

We tested if this system can be used to evoke perceptible activity. To assess this, we first trained water-restricted, head-fixed mice to acquire the Go/No-Go task rule by training them on a simple olfactory discrimination task (Fig. 3A,B). Once proficient, the mice were trained on the detection task. This task was to associate the optogenetic stimulation of mitral and tufted cells with the water reward by generating anticipatory licks selectively on rewarded trials. One group of mice learned to distinguish 5 stimulations from no stimulation (n = 4 mice), and another group learned to distinguish 3 stimulations from no stimulation (n = 12 mice). Both groups of mice learned to detect the presence of stimuli accurately in a few days, although it took slightly longer for the mice to learn with 3 stimulations than with 5 stimulations (Fig. 3C-G; Trials to criterion performance for 3 vs. 5 stimulations = 306.7 ± 1.08 vs. 112.5 ± 50.7; Mean ± SD; p = 0.0056, Wilcoxon rank sum test). We also tested if a single stimulation can be detected in some of these mice (Fig 3F; n = 8 mice). The behavioural performance was well above chance, indicating that the mice perceived a single activation consistently. On the other hand, the behavioural performance was at the chance level in mice where the cannula tips ended up outside of the brain or when the patch cable transmitting the blue light from the source was decoupled from the implanted cannula (Supplemental Fig. 4). These results suggest that targeted optogenetic activation was the salient cue and that the mice readily detect the optogenetically evoked stimulation of mitral and tufted cells.

**Figure 4:**
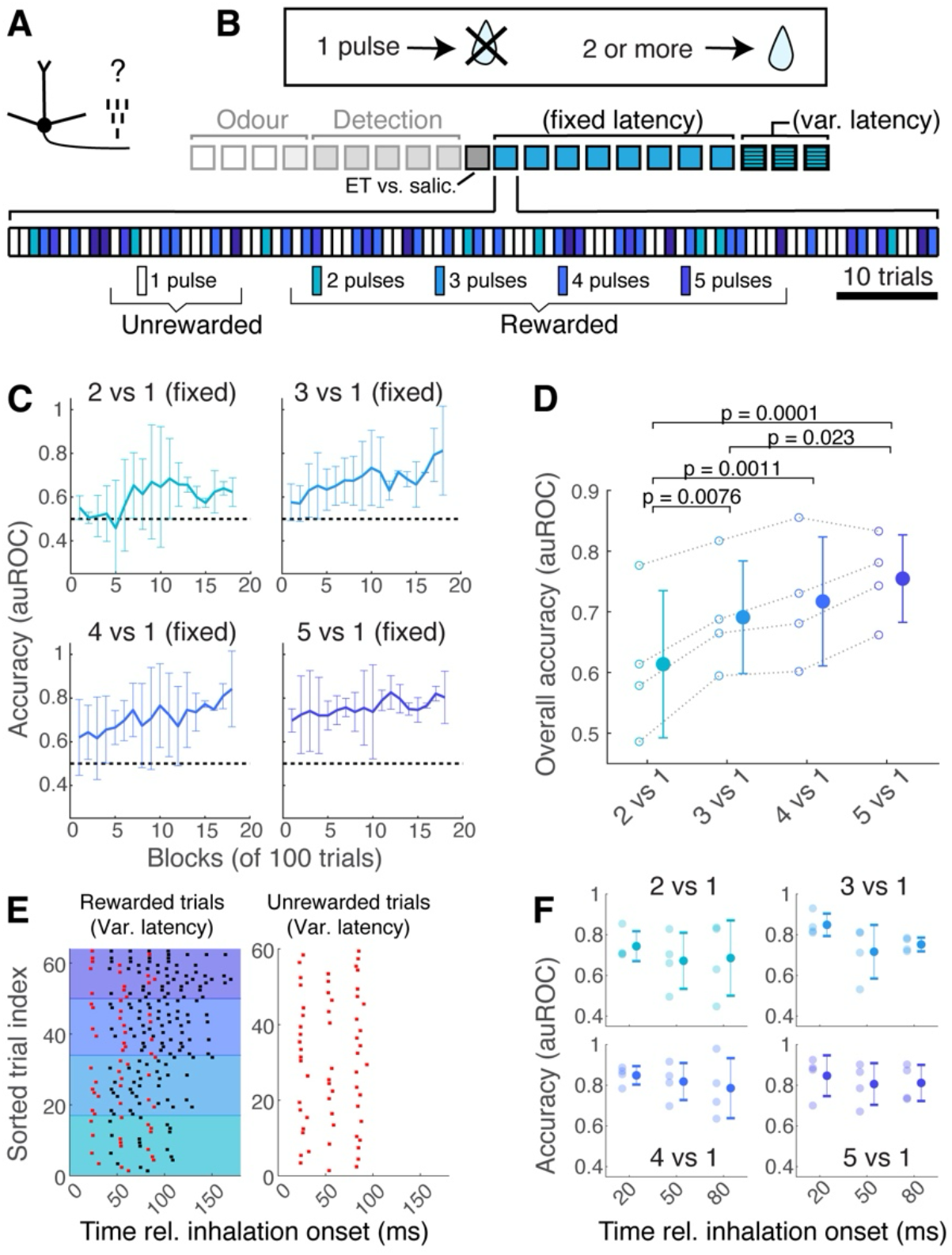
Mice perceive distinct stimulation numbers of olfactory bulb output. **A**. Hypothesis tested: is the number of mitral and tufted cell stimulations distinguishable behaviourally? **B**. Task rule. The unrewarded stimulus was a single stimulation of the LOT. Rewarded stimulus comprised 2, 3, 4, or 5 stimulations of the LOT. These stimuli were semi-randomly chosen. 50% of the trials in a given session were unrewarded trials. Mice underwent this task training after successfully learning to detect light. Timing of light presentation was fixed relative to the sniff cycle. **C**. Learning curves are broken down by the number of stimuli used. N = 4 mice. **D**. Summary data showing the average accuracy across all days. P-Values are from multiple comparisons following a 2-way ANOVA test. **E**. Raster plot showing the timings of light pulses used when the onset latency was varied with respect to the inhalation onset. The timing of the first light pulse is shown in red; the following pulses are shown in black. Left panel = rewarded trials, sorted by the number of pulses used. Right panel = unrewarded trials. **F**. Behavioural accuracy broken separated by the onset of the light pulse. Each panel shows the discriminability of 1 stimulation vs. 2, 3, 4, or 5 stimulations. N = 4 mice. Mean and SD shown.

### Mice perceive differences in the evoked spike counts

Having demonstrated that the evoked activity is perceptible, we asked if the firing rate was a distinguishable feature of the olfactory bulb output. We designed a new Go/No-Go paradigm (Fig. 4A,B), where mice were trained to distinguish different numbers of optogenetic stimulations (1-5). Trials with 1 light pulse were unrewarded, whereas trials with 2, 3, 4, or 5 light pulses were rewarded. Stimulus presentations were semi-randomised (see Method). Since the mice had already learned to detect the presence of optogenetic stimuli (3 stimulations), the mice showed above-chance performance from the beginning when distinguishing 5 pulses against 1 pulse (Fig 4C,D; mean accuracy for the first session = 0.75 ± 0.072, auROC; Mean ± SD; n = 4 mice). The accuracy was lower for the fewer pulse numbers but steadily increased throughout the sessions. Distinguishing 2 stimulations against 1 stimulation was at the lowest accuracy, but the accuracy was above chance after several training days (mean accuracy during the last session = 0.61 ± 0.12, auROC; Mean ± SD; n = 4 mice). Overall, there was a clear dependence on the number of pulses (p = 20.7×10^−12^, 2-way ANOVA; n = 4 mice). The result indicates that the perception of different numbers of evoked action potentials is distinct, where a single pulse difference is perceptible.

Does the ability to distinguish the number of evoked action potentials depend on the timing relative to the sniff cycle? To address this, the onset of the first light pulse was set to one of three latencies (Fig. 4E; 20, 50, 80 ms) with respect to the inhalation onset, which was randomly chosen for a given trial. The ability to distinguish the number of stimulations did not significantly depend on the latencies tested (Fig. 4F; p = 0.12, F = 2.25; effect of latencies in 2-way ANOVA; n = 4 mice), indicating that timing within the respiratory cycle may not matter. Altogether, the results demonstrate that the numbers of evoked action potentials are discriminable, but the ability to distinguish them is not significantly affected by the timing with respect to the sniff cycle.

### The timing of olfactory bulb output activations relative to the sniff rhythm is not perceptible

To further study the importance of sniff timing, we designed a more direct test where the task relied on paying attention to the timing. Mice were trained to discriminate between distinct timing of activations within the sniff cycle (Fig. 5A,B). The rewarded stimulus comprised 3 pulses of light with an onset latency of 20 ms after the inhalation onset. The unrewarded stimulus comprised 3 pulses of light with the onset latency at 120 ms. These stimuli corresponded to mitral and tufted cell activations during the inhalation phase vs exhalation phase of the respiratory cycle, respectively (Fig. 5C). None of the mice exhibited any sign of learning, even with more than 1000 trials of training (Fig. 5D; mean predictor value (an increase in the odds with trials) = −0.0012 ± 0.0021; mean ± SD; n = 3 mice). To rule out that this result is specific to the choice of latencies used, we also trained the mice to discriminate between 20 ms vs 80 ms, followed by 80 ms vs 130 ms (Fig. 5E). The behavioural accuracy remained at the chance level for all conditions used (Accuracy in the last session (auROC) = 0.463 ± 0.086, 0.512 ± 0.023, and 0.545 ± 0.060 for 20 ms vs 80 ms, 20 ms vs 120 ms, and 80 ms vs 130 ms, respectively; n = 4, 3, and 4 mice; mean ± SD).

**Figure 5:**
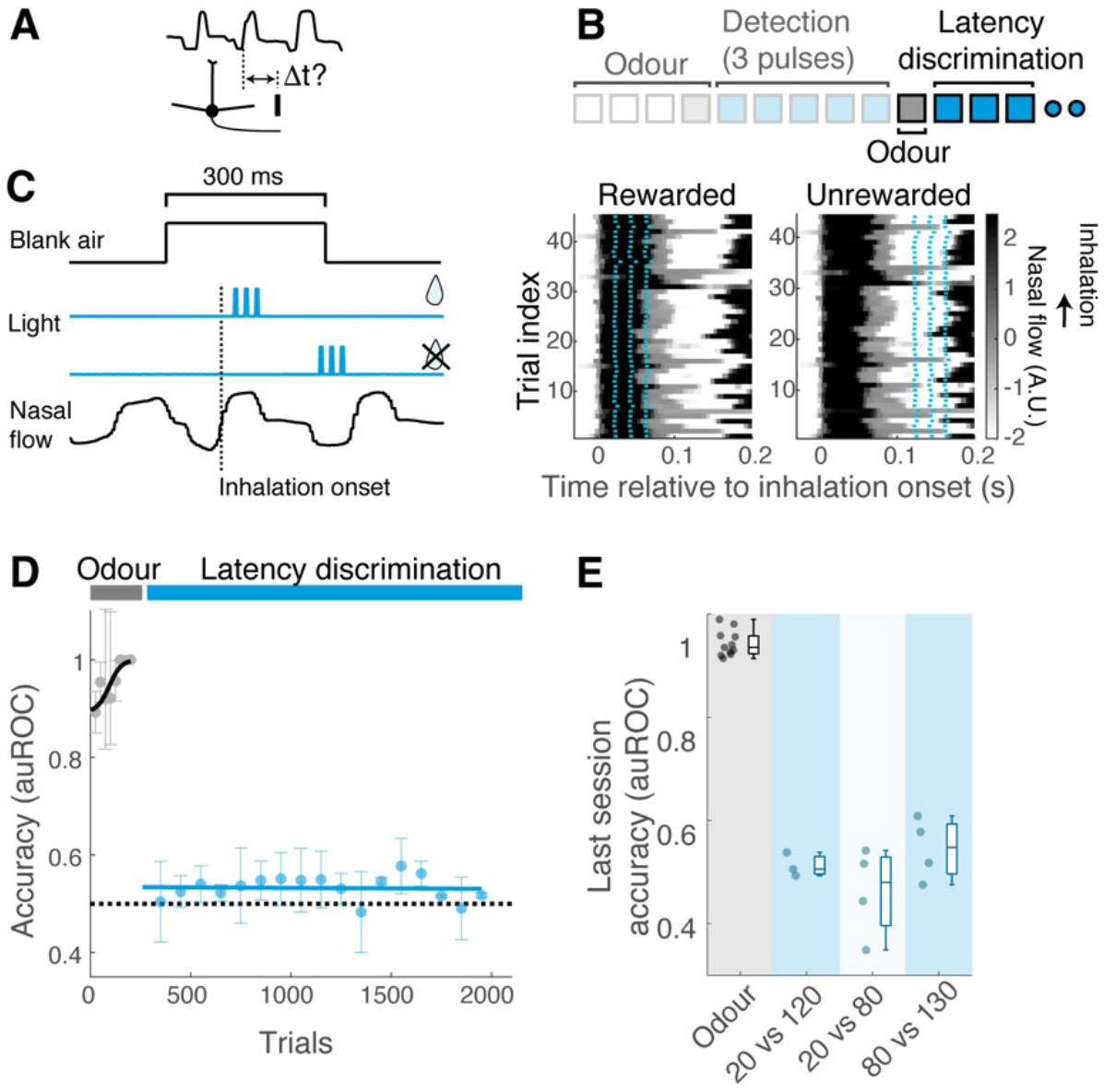
Timing with respect to the sniff cycle is not distinguishable. **A**. Hypothesis tested: Is the timing of mitral and tufted activations relative to the sniff cycle perceptible? **B**. Experimental timeline. Mice underwent the latency discrimination task after acquiring the light detection task. Mice were trained in at least 3 sessions. **C**. Left. Task rule and trial structure. Rewarded and unrewarded stimuli were both 3 light pulses (50 Hz, 5 ms duration each) but differed in the timing with respect to the inhalation onset. Right. Raster plot showing the timings of light stimulations (light blue ticks) for an example session, where rewarded vs. unrewarded stimuli had latencies of 20 ms vs. 120 ms relative to the inhalation onset. Colormap shows the flow sensor reading, where darker colour indicates inhalation. **D**. Learning curve for the odour discrimination (black) and latency discrimination (20 ms vs. 120 ms; light blue). N = 3 mice. Mean and SD shown. **E**. Summary of behavioural accuracy for the last session of training for the 3 pairs of latencies tested (blue) compared to the accuracy of discriminating odours just before. 4 mice were trained on 20 ms vs. 80 ms discrimination followed by 80 ms vs. 130 ms discrimination. 3 mice were trained on 20 ms vs. 120 ms discrimination.

The behavioural insensitivity to the timing of evoked activity was unexpected, given that the olfactory bulb neurons lock prominently to the sniff rhythm. We hypothesised that the optogenetic stimuli may have been too strong in the above experiment, such that they overrode a subtler, baseline sniff-locked activity that the network may use as a reference to compute the spike latencies. If this were true, using weak optogenetic stimulations may be needed to reveal the behavioural sensitivity to this timing. To address this, we used a ramp profile to reduce synchrony in the evoked activity (Supplemental Fig. 5A-E). In addition, the overall light irradiation was lowered to the level that is just detectable (Supplemental Fig. 5F). However, even with this minimal stimulation approach, the mice were not able to distinguish the evoked timing with respect to the sniff cycle (Supplemental Fig. 5G). Based on the above results together, we conclude that the timing of olfactory bulb activations relative to the sniff rhythm is not a perceptible feature.

### Synchronous activations of olfactory bulb outputs are perceptually distinct from asynchronous activations

Does timing matter at all for the downstream targets of mitral and tufted cells? Another aspect of sniff-dependent olfactory encoding is the relative latencies between the olfactory bulb outputs (Fig. 6A). Such latencies, for example, may represent the relative tuning of receptors associated with the ligand presented, where early responses corresponding to greater tuning ^21^.

**Figure 6:**
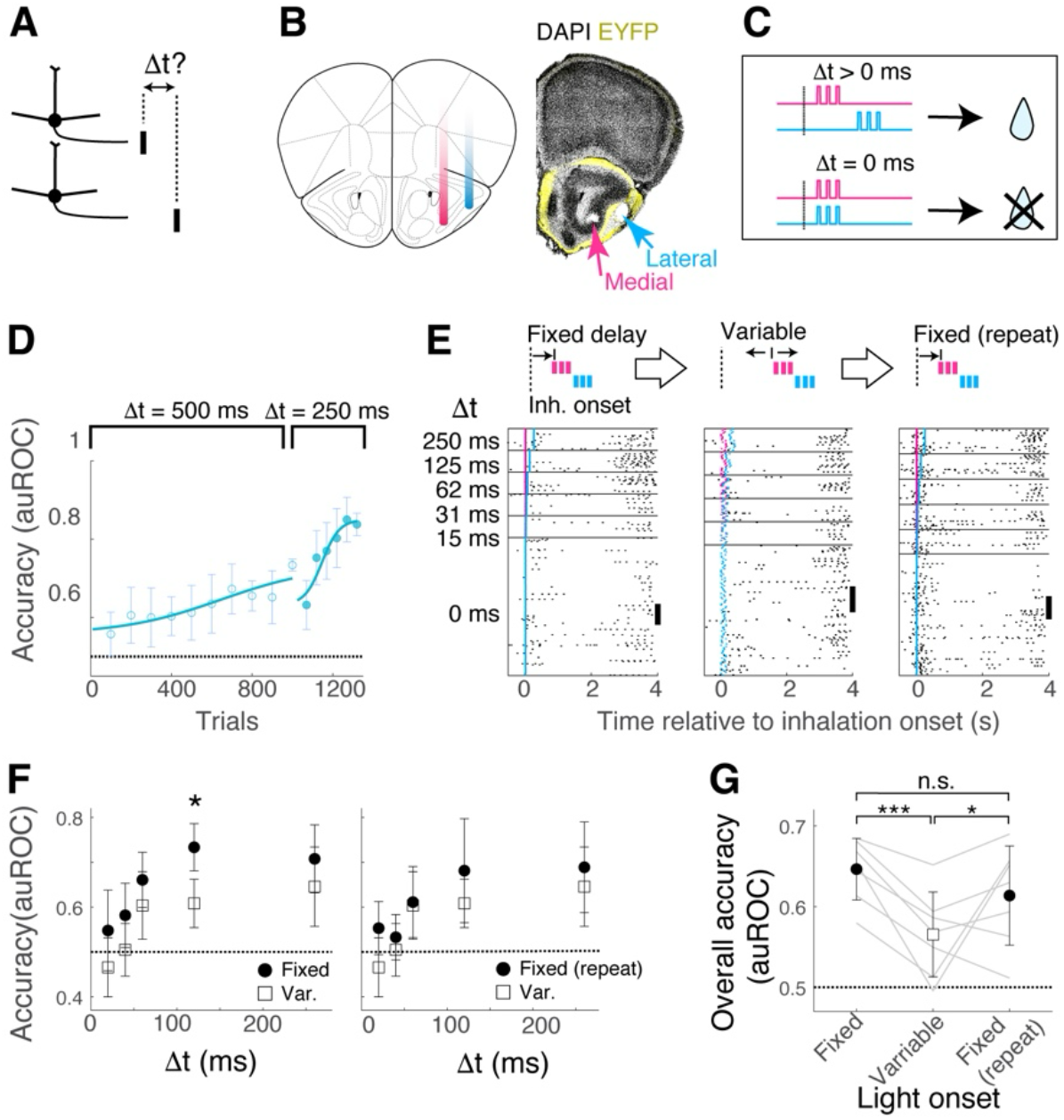
Relative timing of mitral and tufted cell activations is perceptible. **A**. Hypothesis tested: is the relative timing of the olfactory bulb output a perceptible feature? **B**. An experimental approach to recruit two sets of mitral and tufted cells. A dual-core cannula was implanted to target the lateral vs. medial portions of the lateral olfactory tract. **C**. Task structure. The unrewarded stimulus was synchronous stimulations of axons at the lateral and medial locations. The rewarded stimulus was asynchronous stimulations. Dt corresponds to the latency difference between the lateral vs. medial stimulations. **D**. Learning curves for ⍰t = 500 ms, followed by ⍰t = 250 ms for the rewarded stimulus. **E**. Once the mice were proficient at ⍰t 250 ms, different values of ⍰t for the rewarded trials were explored within the same session. Further, the onset relative to the inhalation start was fixed for the first session (left panel), then made variable the next session (middle panel) and returned to the fixed condition on the last day (right panel). Lick raster plots are shown for the corresponding trial types. Scale bar = 20 trials. **F**. Accuracy of distinguishing asynchronous vs. synchronous stimulations for the different ⍰t. The accuracy for the sessions with fixed latency vs. variable latency relative to the sniff cycle (filled vs. hollow symbols, respectively) are shown. **G**. Overall accuracy (all ⍰t combined) for the stimulations with fixed and variable latencies. P = 6.81 × 10-5, 0.169, and 0.026 for fixed vs. variable onset, fixed (repeat) vs variable onset, and fixed vs. fixed (repeat), based on multiple comparisons following a 1-way ANOVA test. N = 7 mice. Mean and SD shown.

To address if relative timing between the output neurons is perceptible, we used another paradigm that trains mice to distinguish synchronous vs asynchronous activations of olfactory bulb outputs. This paradigm is based on a previous study by Rebello et al ^41^, except that the axons of mitral and tufted were stimulated instead of their cell bodies. We implanted a dual cannula where two optical fibres targeted the lateral vs. medial portions of the lateral olfactory tract (Fig. 6B,C). Trials with simultaneous stimulations of the lateral and medial locations were unrewarded, while trials with staggered stimulations were rewarded (Fig. 6C). The onset of the first light pulse was fixed to 20 ms from the inhalation onset. We first confirmed that the mice can detect the optogenetic stimulations at both locations of the lateral olfactory tract. Then, the mice were trained to distinguish the 500 ms latency difference from the synchronous stimulation. This training was followed by a shorter latency difference for the rewarded trials (250 ms). Mice were able to acquire this task (Fig. 6D; mean number of trials to criterion performance = 600.8 ± 139.4 from the start of 500 ms training; mean ± SD; n = 5 mice). Once proficient, we explored the temporal resolution of the task performance by presenting a range of latency differences (15 ms, 31 ms, 62 ms, 125 ms, and 250 ms). The accuracy of distinguishing between synchronous and asynchronous stimulations depended on the interval (Fig. 6E,F; p = 8.61 × 10-5, F = 8.71, 1-way ANOVA; mean accuracy = 0.55, 0.58, 0.66, 0.73, 0.71 in increasing interval duration; n = 7 mice).

A previous report suggested that the temporal resolution of this task is better if the stimuli arrive at a consistent phase of the sniff cycle ^41^. We tested this by presenting our light stimuli with a fixed delay relative to the sniff cycle or with variable timing. To allow within-animal comparison, the mice underwent fixed vs. variable timing conditions on alternate days over three sessions (“Fixed” − “Variable” – “Fixed-repeat”). This comparison revealed a slight tendency for the behavioural performance to be more accurate when the stimuli were delivered at a fixed time latency from the inhalation generally (Fig. 6F, G; p = 7.26 × 10^−5^, F = 10.09, the effect of trigger condition in 2-way ANOVA), indicating that the sniff rhythm may modulate the state of downstream regions. Overall, we conclude that the relative timing of action potentials between neurons is perceptible (Fig.7). While the respiratory cycles may not serve as a reliable reference timing signal, the sniff-locked rhythmic activity may still exert a modulatory effect on the quality of discriminable patterns of activity.

**Figure 7:**
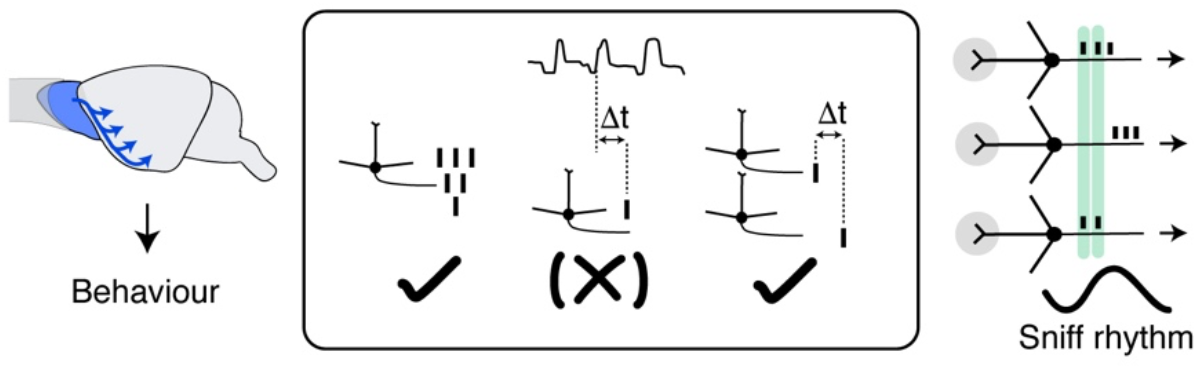
Schematic of the finding. The firing rate and synchrony, but not the latency relative to the sniff cycle, are perceptible features of activation patterns in the temporal domain. Timing with respect to the sniff cycle, however, modulates the quality of evoked patterns. Synchronous arrival of stimuli coupled to an animal’s sniffing may temporally align the activity of mitral and tufted cells associated with different olfactory receptors.

## Discussion

The olfactory system has attracted much attention as a model system for studying the temporal encoding of stimuli. Sniff-locking is a prominent feature of activity patterns encountered in the early stages of olfactory processing ^20,23,24,26,28,32,34,39,44^. As a result, it has been widely thought of as a fundamental timing mechanism for decoding olfactory information, much like other systems that use oscillations for encoding states ^45^.

Our results show that, despite its prominence, the sniff rhythm – the timing within the sniff cycle itself - is not perceptible downstream of the olfactory bulb. This finding points to a possible difference between the input to the olfactory bulb vs. its output. Input to the olfactory bulb contains mechanosensory signals from the olfactory sensory neurons, which follow the rhythmic air flow from sniffing ^27,44^. The resulting rhythmic glutamatergic input to the olfactory bulb forms the basis of baseline sniff-locking by the olfactory bulb neurons ^20^. This availability of a reliable timing signal may be why the sniff phase is perceptible when the olfactory nerve is stimulated ^22^. However, this common rhythm is subject to local inhibition in the olfactory bulb, which diversifies the baseline sniff-locking of mitral and tufted cells ^28,46^. Therefore, while the latency code requires an internal reference signal ^47^, downstream of the olfactory bulb, a reliable timing signal about sniffs may not be available, making the timing relative to sniffing an unreliable code for olfactory cortices. The behavioural insensitivity to the spike timing relative to the sniff cycles is consistent with a previous work, which reported that once mice learn an optogenetically evoked pattern of activations, the pattern is recognised indiscriminately at all sniff phases ^40^. However, our result showed a slight dependence on the triggering condition for discriminating the synchronous vs. asynchronous activations. Therefore, the sniff rhythm may still play a subtle, modulatory role downstream in refining the quality of unique patterns of activations.

Some aspects of temporal coding in the olfactory bulb are, however, perceptible. With the dual-core experiment, we confirmed the importance of relative timing – in particular, synchrony - between the activity of olfactory bulb output neurons. This result is consistent with previous findings ^40^. Another interpretation of this result is that the downstream regions are tuned to specific intervals. Some pyramidal cells of the piriform cortex are known to be tuned to specific input intervals ^48^. Intervals present in the olfactory bulb output may be synchronised downstream using differential bias currents ^49^, conduction delays ^50^, or well-timed inhibition as in the Reichardt detector ^51^. Be it synchrony or detection of specific intervals, there is a requirement for a common timeframe for the relative timing between neurons to be meaningful. Here, the sniff rhythm likely plays a crucial role in aligning the activity of mitral and tufted cells associated with different olfactory receptor types under a common clock.

Our study also points to the importance of the firing rate in olfactory encoding at a relatively early stage of olfactory processing. This aspect of activity patterns may benefit from greater attention in the future. In our experiments, the downstream targets of mitral and tufted cells include the anterior olfactory nucleus, tenia tecta, olfactory tubercle, piriform cortex, lateral entorhinal cortex, cortical amygdala, medial amygdala ^11,34,52–55^. The decoding mechanism may be unique for each region, where only one aspect of the encoding format is relevant to the specific downstream area. Alternatively, the firing rate and timing are both used to convey different aspects of olfactory stimuli, such as the concentration and identity of odours, and the target structure to possess mechanisms to decode multiplexed signals ^56^.

In general, the specific format used to encode a sensory stimulus is a unique solution for each area and its computation. Precisely timed spikes are integral for encoding time-sensitive stimulus features, as seen widely in the early stages of visual, auditory, mechanosensory, and electro-sensing systems ^51,57–62^. Elsewhere, in theory, the information transmission rate is higher using precise timing ^63,64^, including in conveying changes in the environmental statistics ^3^. In practice, other factors, for example, noise and mechanisms of synaptic integration *in vivo*, may make this coding format unsuitable ^65–67^. We do not know if other systems in the brain that use oscillation-based timing, such as phase advances in the place cell encoding ^45,68,69^, offer a more reliable and behaviourally salient reference signal. Our study reaffirms the importance of direct tests to establish a link between features of activity patterns and perception and redefine the utility of internally generated rhythms to encode the environmental cues that drive a rich repertoire of behaviours.

## Materials and methods

*Animals*. All the animal experiment procedures were approved by the Animal Care and Use Committee (ACUC) of Okinawa Institute of Science and Technology Graduate University [Protocol No. 2023-050]. Adult heterozygous Tbx21-Cre::Ai32 mice were admitted to this study (21 male and 14 female mice, 10 - 17 weeks old; see the methods for Light Detection task for the inclusion criteria). Mice were generated by crossing Tbx21-Cre (Haddad et al., 2013) mice with Ai32 (Madisen et al., 2012) mice. Tbx21-Cre (stock numbers: 024507) and Ai32 (stock numbers: 012569) mice were originally purchased from the Jackson Laboratory.

### Surgery

Animals were deeply anaesthetised using isoflurane (IsoFlo, Zoetis Japan) and fixed in a stereotaxic frame (#1900, Kopf Instruments). Analgesia was provided by i.p. injection of Carprofen (Rimadyl, Zoetis, Japan) at 5mg/kg body weight, and body temperature was maintained at 33-35 °C (#40-90-8D, FHC USA). The surgical field was shaved and disinfected using iodine (Mundipharma Japan). The skin was resected, a fibre cannula or cranial window was implanted, and a metal head plate was glued using a cyanoacrylate gel (Loctite, Germany) posterior to the bregma. The head plate was then secured by dental acrylic (Kulzer, Hanau, Germany). All animals were monitored daily and administered carprofen for 3 consecutive days post-surgery and allowed to recover at least 2 weeks before the start of behavioural experiments.

### Details of optical fibre implantation

Bregma and Lamda, visible on the exposed skull, were used to adjust the anterior-posterior angle such that the height difference between the two landmarks was zero. The mediolateral axis was adjusted by making the skull height at −1.5mm/±1.5mm AP/ML relative to bregma zero. For single-core fibre implantation, a unilateral craniotomy on the left side of the brain was made at AP 2.1 mm, ML −1.6 mm from bregma. A 0.39NA, 200um diameter fibre (CFMC12L05 Thorlabs, USA) was inserted 4.71mm at a 28.5-degree angle from the centre. For dual-core fiber implantation, a unilateral craniotomy was made at the coordinate +0.2mm/0.9mm (AP/ML relative to Bregma). The two fibres (#CFM32L10, Thorlabs, USA) were implanted with the medial fibre insertion at +0.2mm/0.9mm (AP/ML relative to Bregma) with a pitch of 30° relative to the dorsoventral axis and an additional rotation of 30° around the fibre’s central axis to make the lateral core targeting more anterior. Fibers were advanced 4.75mm relative to the brain surface. In both cases, the fibre was secured with dental acrylic.

### OB craniotomy window implantation

A cranial window of ∼1.5 × 1 mm was implanted on the left olfactory bulb by replacing a piece of skull with a hand-cut glass coverslip (thickness No. 1; Matsunami, Japan). The window was glued with cyanoacrylate (Histoacryl, B. Braun, Germany) and dental acrylic. A plastic tube cut from a dust cap (CAFP, Thorlabs, USA) was implanted above the cranial window, which served as an adapter for the optic fibre. Dental acrylic with black nail power (Diamond Acrylic Powder, Glam and Glits, South Africa) mixed was mounted around the cap to prevent the light from leaking. The open end of the implanted cap was sealed by a hand-cut silicon plug to prevent dust from accumulating in the tube. The cranial window was cleaned when the plug detached.

### Olfactometry

Odours were presented as described previously ^70^, except for the odour pulse duration (0.3 seconds). Odour concentrations were approximately 1% of saturated vapour with an average mass flow rate of 2 lpm including the carrier air. Odorants were purchased from Sigma-Aldrich (Ethyl butyrate, W242705; Ethyl valerate, 290866; Butyl acetate, 287725) and Tokyo Chemical Industry (Methyl tiglate, T0248; Methyl valerate, S0015; Methyl salicylate, V0005; Eugenol, A0232, Ethyl tiglate, T0247; Butyl butyrate, B0757; Methyl anthranilate, A0500; Acetophenone, A0061; Salicylaldehyde, S0004; Methyl butyrate, B0763). The odorants were stored in a dark, air-tight cabinet filled with nitrogen.

### Behaviour – Go/Nogo discrimination

Water-restricted mice were first habituated to head fixation and licking on a running wheel for 3-5 days. Lick responses were measured using an IR beam sensor (PM-F25, Panasonic, Japan). Once habituated, mice were trained to learn the association between one odour (ethyl butyrate) with water reward, as well as between another odour (methyl tiglate) and a lack of reward with methyl tiglate by generating anticipatory licks selectively for the rewarded stimulus. This initial training took place until the mice reached the criterion level of accuracy (auROC = 0.8). This took 2-4 sessions, and one session took place per day. Each trial began with an LED cue (single pulse blue light at 11 mW for 640 ms presented ∼4 s before the odour valve onset). Blue LED in the chamber was turned on simultaneously with the final valve opening for all trials, to accustom the mice to blue light presentations. The trial type (rewarded or not) is generated semi-randomly, avoiding more than three consecutive trials of the same reward condition. Water rewards were given unconditionally 4.2 seconds after the FV opening (20 ul, delivered over 2 droplets). Once proficient, mice were trained on other pure odor pairs or optogenetic stimuli. For all Go/No-Go paradigms, the trial order was semi randomised, such that, if the 3 previous trials had the same reward contingency, the trial type for the following trial was forced to be different. Otherwise, the trial type was shuffled. Before each optogenetic behavioural experiment, the mice learned to discriminate a new pair of odours.

### Optogenetics with behaviour

A simple closed-loop system was implemented by running parallel scripts, first for online analysis of sniff signals and second for controlling the behavioural experiment and generating light pulses; sniff patterns of head-fixed mice were monitored continuously by placing a flow sensor (AWM3100V, Honeywell, USA) adjacent the right nostril. The signal was acquired (NI USB 6210 DAQ, National Instruments, USA) at 1 kHz. Inhalation onsets were detected by monitoring for a positive threshold crossing online and binarised to a TTL signal using a LabVIEW script (inhalation onset time). The digital signals corresponding to the inhalation onset times are sent to a separate script that controls the behavioural experiment. The trial structure of the behavioural experiment was similar to that of the olfactory discrimination task, except that in place of the odours, specific light stimuli were delivered through the implanted cannulae. The final valve was still opened to present blank air to mimic the olfactory behavioural experiment as much as possible—the first inhalation onset after this valve opening was used to trigger the optogenetic stimulus delivery.

Commands to the LED driver were generated using a custom-written code using waveform functions of LabVIEW embedded in the behavioural control script (temporal resolution = 1kHz) and outputted using an analogue module (NI USB-6531, National Instruments, USA) controlled by a custom LabVIEW script. These were used to generate light outputs using a 450-nm laser diode (L450P1600MM, Thorlabs, USA) driven by a T-cube LED driver (LEDD1B, Thorlabs, USA) for single-fibre experiments or a 4-channel LED driver (DC4104, Thorlabs, USA) for dual-fibre experiments.

The light from the laser diode was collimated and then delivered to the patch cable. We used a 0.39NA, 200um patch cable (MR77L01, Thorlabs, USA) for the single-core LOT experiment and OB window experiment, and a 0.39NA, 200um dual-core patch cable (BFY32SL1, Thorlabs, USA) for dual-core LOT experiment. To deliver the light through the implanted fibres, the patch cable was connected to the implanted cannula by a ceramic mating sleeve (ADAF1, Thorlabs, USA). For the weak olfactory bulb stimulation, the light was presented through an implanted window by attaching the patch cable to an adapter that had been implanted just above the cranial window. The gap between the patch cable ending and the cranial window was approximately 1.5mm to diverge the beam area to ∼0.4 mm^2^ at the brain surface. The light output was calibrated daily using a power meter (PM320E, Thorlabs, USA). Unless otherwise stated, the onset of the first light pulse was set to 20ms after the inhalation onset.

### Light detection task

The rewarded stimulus was the presence of light (triplets at 50 Hz, 25 mW, 5-ms duration, presented 20ms after the inhalation onset). The unrewarded stimulus was no light. Only the mice that reached the criterion level in this task (auROC = 0.8) were included in further experiments. At the end of training, 4 mice participated in control experiments, where the patch cable was uncoupled so that the light output from the patch cable was pulled away from the implanted cannula. In addition, data from further 5 mice that failed to learn the detection task, where the cannulae were confirmed to be outside of the brain upon post-hoc, are shown in the Supplemental Fig. 4).

### Pulse number discrimination task

Rewarded stimuli were 2, 3, 4, or 5 pulses of light (25 mW, 50 Hz, 5 ms duration each), presented with the frequency ratio (1:3:1:1, respectively). The unrewarded stimulus was 1 pulse of light (25 mW, 50 Hz, 5 ms duration). Training contained two stages. In the first stage, stimuli were presented at fixed latencies relative to inhalation. In the second stage, stimuli were set to one of the following latencies relative to inhalation: 20 ms, 50 ms, or 80 ms.

### Latency discrimination task

Mice were trained to lick for the light stimuli presented at a shorter latency after the inhalation onset, and refrain from licking for stimuli that occurred at a later latency.

### Synchronous vs. asynchronous activations

The rewarded stimuli were asynchronous optogenetic stimulations using the lateral and medial fibre cores. The unrewarded stimulus was the synchronous stimulations. Initially, the interval for the rewarded stimulus was 500 ms (the lateral core followed the medial core). Then, the interval for the rewarded stimulus was reduced to 250ms. After that, the rewarded stimuli consisted of a set of intervals [15 ms, 31 ms, 62 ms, 125 ms, 250 ms]. The stimuli were either triggered at a fixed latency (20 ms) after inhalation onset for the “Fixed delay” experiments, and triggered with a random latency in the range of 0-200 ms for the “Variable delay” experiment.

### In vivo electrophysiology – acute recording from anaesthetised mice

4 mice that had previously been used for the behavioural experiments (optogenetic detection and latency discrimination) were used but had access to water *ad libitum* for at least one day before the recording. Mice were anaesthetised with isoflurane (3% for induction, 1% for maintenance), and a small craniotomy was made over the left olfactory bulb. After the craniotomy was done on the left olfactory bulb, isoflurane was removed, and mice were anaesthetised with (ketamine/xylazine, 100 mg/kg and 20 mg/kg, respectively). Probes were coated with DiI (V22885, Invitrogen) so that the tract in the olfactory bulb was recorded. Extracellular recording of mitral/tufted cells was obtained with a 16- or 32-channel silicon probe (A1×16-Poly 2-5 mm-50s-177, catalog #CM16LP, or A1×32_Poly3_5mm_25s_177, NeuroNexus). Data was acquired at 20 kHz with a low-noise amplifier (RHD2132, Intan Technologies) and RHD controller (C3004, Intan Technologies).

### In vivo electrophysiology – acute recording from awake mice

Electrophysiology for the olfactory bulb stimulations was conducted for one session on naïve, awake animals using the 32-channel probe. Mice were habituated for head-fixation for 3-4 days after a 2-week recovery from head plate implantation. Once habituated, a craniotomy was done on the left olfactory bulb under isoflurane. Isoflurane was removed, and the recording was done on the head-fixed mouse on the treadmill when the animal woke up. The craniotomy was covered by a thin layer of 4% agarose (A9539, Sigma-Aldrich) in Ringer solution for stability. While the probe was in the brain, the light was presented from a fibre optic cannula placed near the craniotomy. The stimulation was randomly associated with a water reward to keep the mice engaged. Electrophysiology using the Neuropixels probe (Neuropixels 1.0, IMEC) was conducted for one session on naive, awake animals as described above, except that odours were presented instead of optogenetic stimuli. The Neuropixels probe was connected to the PCIe card (PXIE_1000 on a PXIe-1082DC chassis, National Instruments) via a headstage (HS_1000). Data from 250 active sites (1:250) was acquired at 30 kHz using the OpenEphys GUI ^71^. Internal tip reference was used. The probe was lowered to 2.5 mm from the dorsolateral surface of the left olfactory bulb.

### Post-hoc histology

After the electrophysiology or the completion of the behavioural experiment, mice were deeply anaesthetised with isoflurane using a mask. Immediately after the cessation of breathing, mice were perfused transcranially first with phosphate buffer (NaH_2_PO_4_(225.7 mM), Na_2_HPO_4_ (774.0 mM), pH adjusted to 7.4) and then fixed with the formaldehyde solution (4% in phosphate buffer). For mice implanted with optic fibre, the fibre was removed 24-72 hours after post-fixation to preserve the fibre tract anatomically. The brain was dissected and sliced to 100μm thickness coronal sections using a vibratome (5100 mz-Plus, Campden Instruments, United Kingdom). Sections were then stained with DAPI (D9542, Sigma-Aldrich). Confocal images were acquired on the Zeiss confocal microscope (Zeiss LSM780, LSM880, or LSM900) with a 10× objective for the cortical sections and a 20× objective for the OB.

### Deviation from intended latency

The flow sensor record was analysed offline in *Spike2* software to detect the onset of inhalation onset. The signal was first conditioned to remove the slow variations in the offset, using the built-in function *DC Remove* with the time constant set to 0.4 s. Inhalation peaks were detected, and the inhalation onset was the first positive crossing of a threshold value (0.08V). Latency to the first light pulse was measured from the measured inhalation onset to obtain the observed latency. This was compared against the intended stimulation latency to get the temporal precision. The data presented in Figure 2 is the deviations from individual trials for an example session, and the value quoted is the average standard deviation.

### Behavioural accuracy

The behavioural accuracy was determined by comparing the number of anticipatory licks produced for rewarded trials vs those for unrewarded trials. All licks generated between the onset of the valve opening and before the water delivery (4.2 s after the valve onset) were counted. The discriminability was determined by measuring the area under the receiver-operating-characteristic curve (auROC) using the Matlab function *per curve*. The auROC ranges from 0 - 1, with 1 being perfect discriminability. The chance level is 0.5 for a binary discrimination task.

### Trials to criterion

To interpolate the number of trials needed to reach the criterion performance in the behavioural tasks (auROC = 0.8), the behavioural accuracy over time was fitted using a logistic function using the Matlab function *glmfit* and interpolated using the Matlab function *glmval*. The trials to criterion were the first trials where the interpolated behavioural accuracy was above 0.8.

### Behavioural analysis: Statistics

Unless otherwise stated, the summary statistics reported are mean and standard deviation. 2-way ANOVA (Matlab function *anovan*) was used to analyse the result of two experiments: (1) 1 stimulation vs. 2 or more stimulations and (2) synchronous vs. asynchronous stimulations. For the analysis of 1 stimulation vs. stimulations, the two factors considered were the training stage (block number) and the number of pulses. In analysing the animals’ ability to distinguish synchronous vs. asynchronous stimulations, the two factors considered were the interval (Δt) and the timing relative to the inhalation onset (‘Fixed’, ‘Variable’, and ‘Fixed repeat’). Multiple comparison test using the Tukey-Kramer method was carried out using the Matlab function *multcompare*.

### Spike sorting

Data were high-pass filtered, and units were sorted offline using Kilosort 4.0 (Pachitariu et al., 2024), with parameters adapted from the recommended parameters from the authors of Kilosort. The channel map was recreated from the layout of the recording electrode. Then, the automated outputs from Kilosort were manually curated with Phy, an open-source graphical user interface (Rossant et al., 2016). Only those clusters with a clear refractory period and waveforms distinguishable from the background were labelled as good single units and used for further analysis.

### Multi-border SVM for decoding the odour ID

We chose a multi-class support vector machine to determine if a specific feature of the population activity from simultaneously recorded mitral and tufted cells contained information to decode which of the 8 olfactory stimuli was presented for the given trial. On average, the number of units recorded per session was 63.8 ± 43.5 (N = 5 mice). For training, Matlab function fitcecoc was used with the prior parameter set to uniform. The model was trained on 90% of trials from a given session, and the accuracy of the classifier was tested using the remaining 10% of trials. On average, these corresponded to 84.2 ± 6.1 trials for training and 9.6 ± 0.5 trials for testing. The trial selection was randomised using the Matlab function datasample and repeated 100 times. SVM based on the firing rate: the number of spikes observed within the temporal window was counted for each unit. Time zero was the start of the odour presentation. The window size was systematically varied, as shown in Figure 1. SVM on latencies: for each unit, the time taken for two action potentials to be generated since the onset of the first inhalation after the odour presentation was extracted.

## Acknowledgements

We thank Yu-Pei Huang and M. Inês Ribeiro for technical assistance, the members of Animal Resource Section, Imaging Section, Mechanical Engineering Section of OIST’s Core Facilities, Sayori Gordon for the administrative assistance, Josefine Reuschenbach, and M. Inês Ribeiro, for helpful comments on the manuscript. This work was supported by OIST Graduate University and Japan Society for the Promotion of Science Grants-in-Aid for Scientific Research (C) 24K09688.

## Supplemental figures

**Supplemental figure 1:**
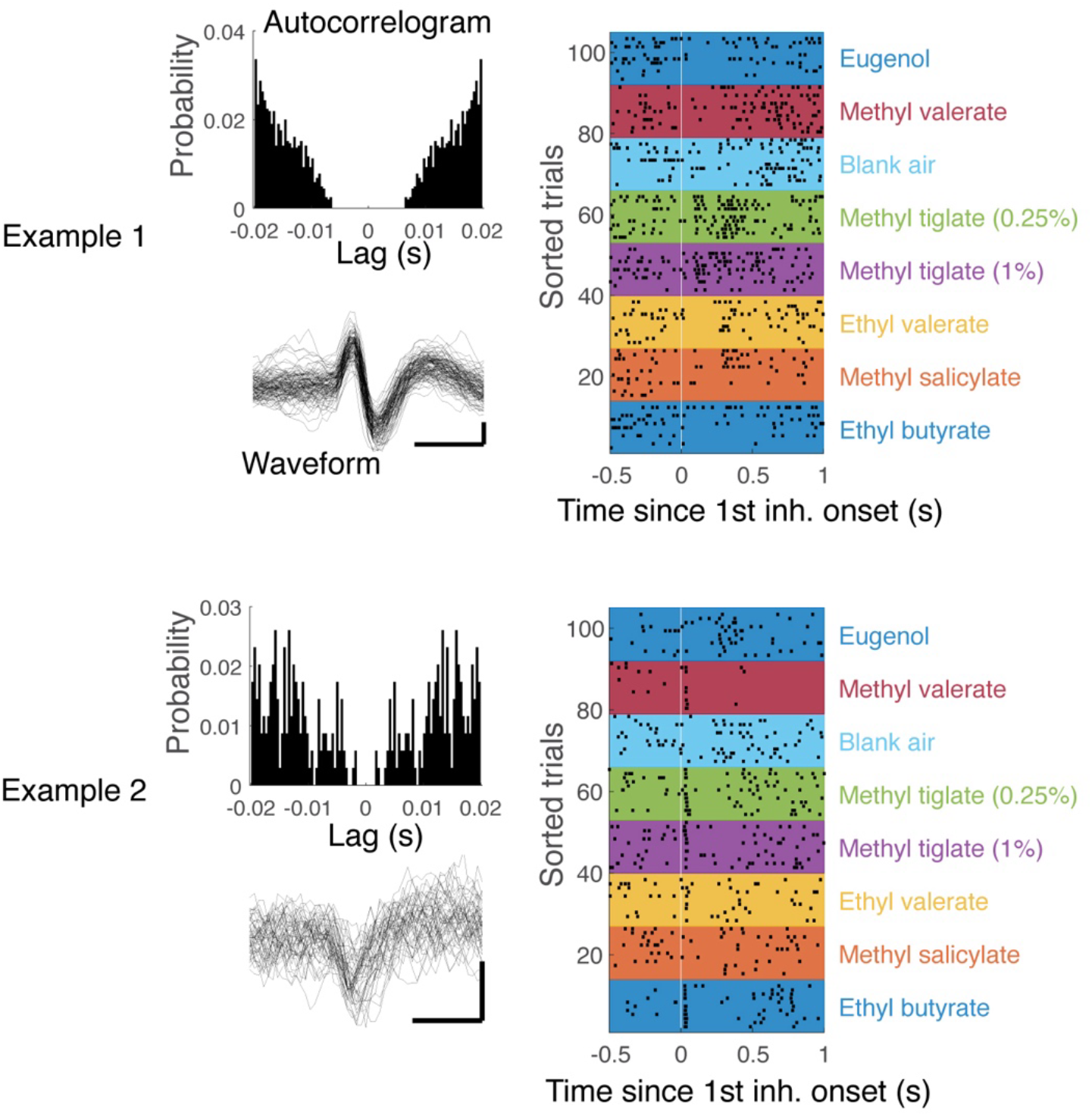
Stability of extracellular recording in awake, head-fixed mice using Neuropixels 1.0. Left. The auto-correlogram and selected raw spike waveforms are shown to demonstrate the recording qualities of two example units for the recording session shown in Figure 1 (top and bottom). Right. Spike raster plots for the example units, showing the timing of spikes observed relative to the onset of first inhalation after the odour onset. Trials are sorted by the odours presented.

**Supplemental figure 2:**
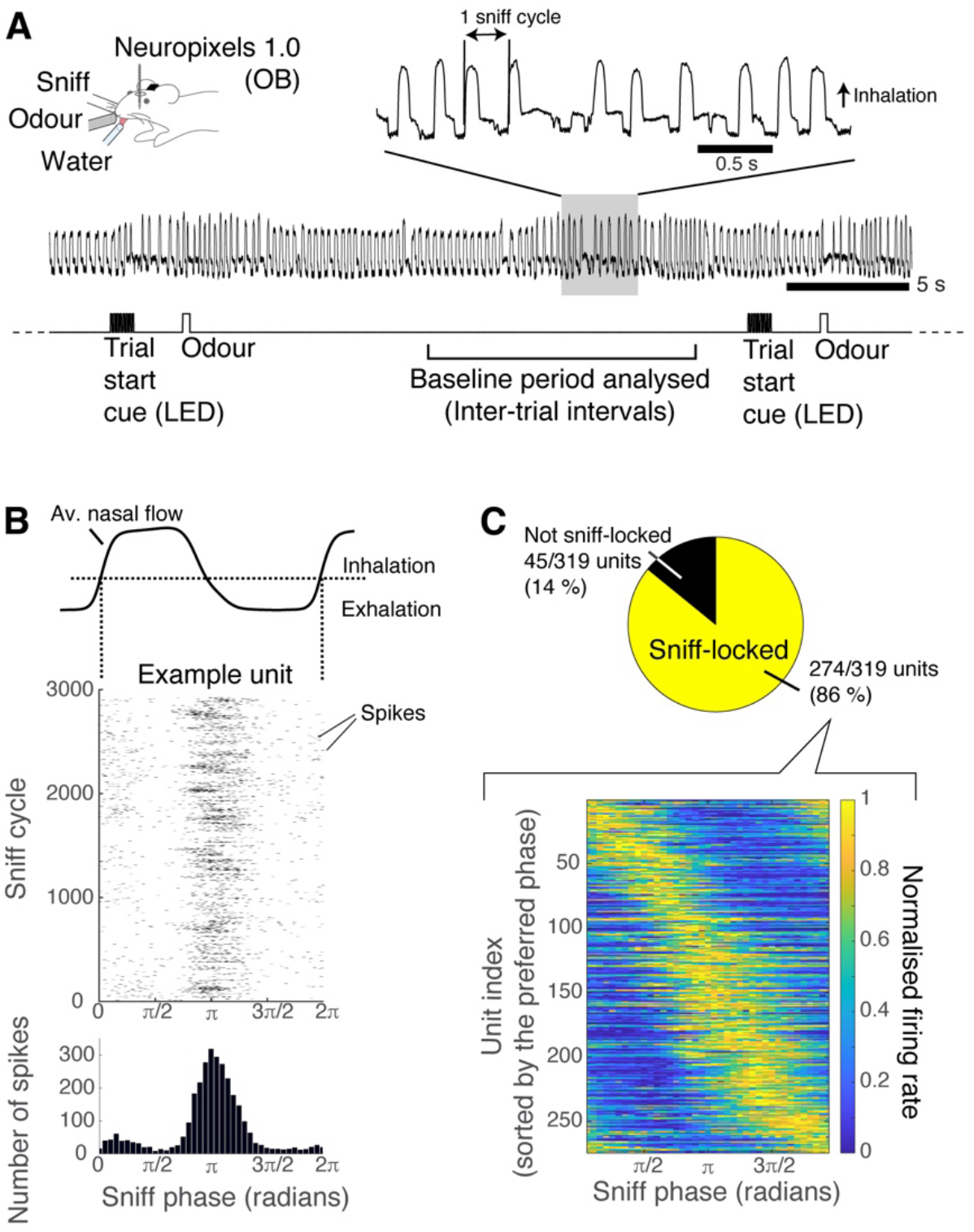
Ample sniV-locking during the baseline periods in awake, engaged mice. **A**. Spontaneous activity from Neuropixels recordings in awake, engaged, and naïve mice was analysed. Inter-trial intervals from the odour exposure experiment were used. The analysed period was from 10 seconds after the previous odour presentation to 5 seconds before the next odour presentation. The sniff cycle was defined from one inhalation onset to the next inhalation onset. **B**. Spike timings from each unit were aligned to the sniff cycles by expressing the timings in terms of phases with respect to the cycle. This spike raster plot from an example unit shows a tendency for spikes to occur in the middle of the sniff cycle. **C**. Sniff-coupled units were defined by those whose sniff-aligned modulation is greater than a shuffled control (see^28^). Most of the recorded units were sniff-locked and varied in the preferred phase.

**Supplemental figure 3:**
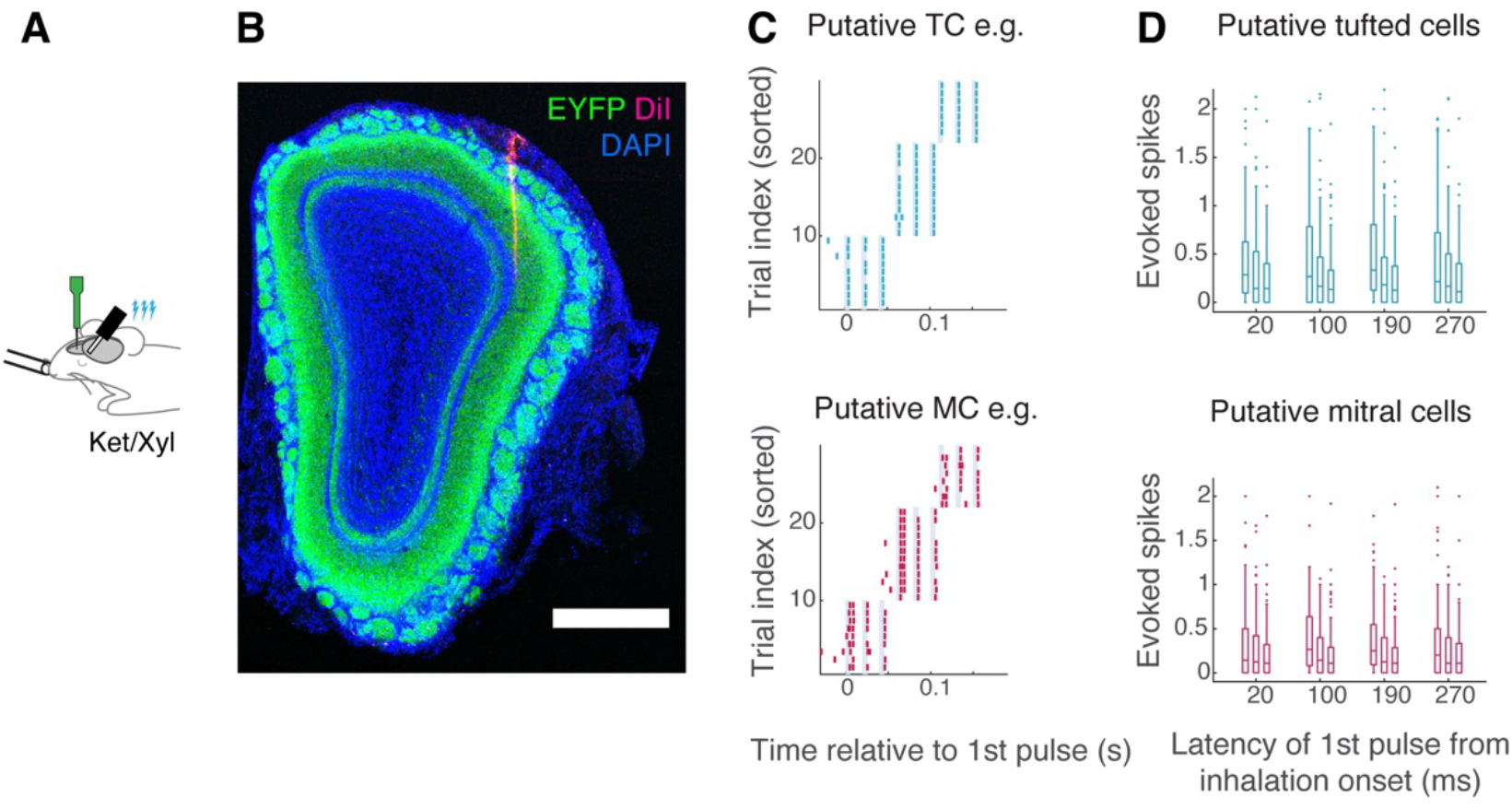
Consistent optogenetic recruitment of mitral and tufted cells across sniV phase. **A**. Experimental configuration: extracellular electrophysiology was conducted in the left olfactory bulb of anaesthetized mice using a 32-channel probe (NeuroNexus). **B**. *post-hoc* histology with DiI showing the electrode track. Scale bar = 0.5 mm. **C**. Units were classified into putative tufted cells and mitral cells based on the phase they lock to at the baseline (Fukunaga et al., 2012). Raster plots for a putative tufted cell (top panel) and a putative mitral cell. Light shading indicates the timing of light presentations (3 rectangular pulses at 50 Hz; 5 ms duration for each pulse; 19.2 mW). Trials are sorted by the latency to first pulse relative to the inhalation onset (onset latencies for the first light pulse used = 20ms, 80ms, and 110ms). **D**. Box plot summary of evoked spikes for the individual pulse within the triplets and separated by the latency to the 1^st^ pulse relative to the inhalation onset. Horizontal lines in the box correspond to the 25^th^, 50^th^, and 7^th^ percentiles, vertical bars indicate the range of the data, and outliers are shown with individual dots. Of the 232 units recorded for this experiment, 89 were classified as putative tufted cells, and 113 were classified as putative mitral cells (n = 3 mice).

**Supplemental figure 4:**
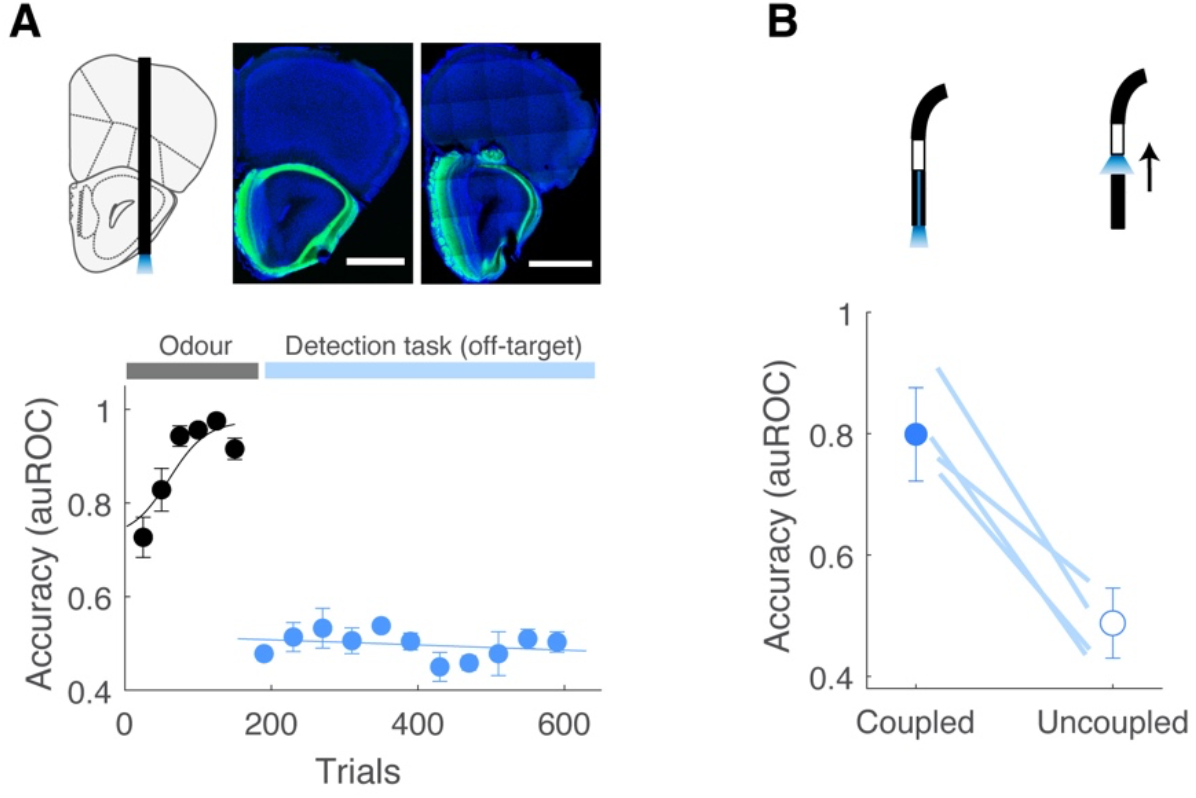
OV-target and uncoupled controls. A) Case 1: optic fibre tip was found to be outside of the brain upon post-hoc histology. Top: Confocal images of a coronal section of the brain from two example mice at the location where cannula tips were located. Scale bar = 1 mm. Bottom: Learning curve for odour discrimination task and light detection task for mice with off-target cannula placement. N = 5 mice. B) Case 2: the patch cable is not coupled with the implanted cannula. Up: The optic cable is inserted to the mating sleeve, but the pin on the cable is not inserted to the alignment pin hole, preventing light from reaching the brain. Down: The accuracy for light detection task when the patch cable and the implanted cannula was coupled (left, blue) and not coupled (right, white) for mice in case 2. N = 4 mice.

**Supplemental figure 5:**
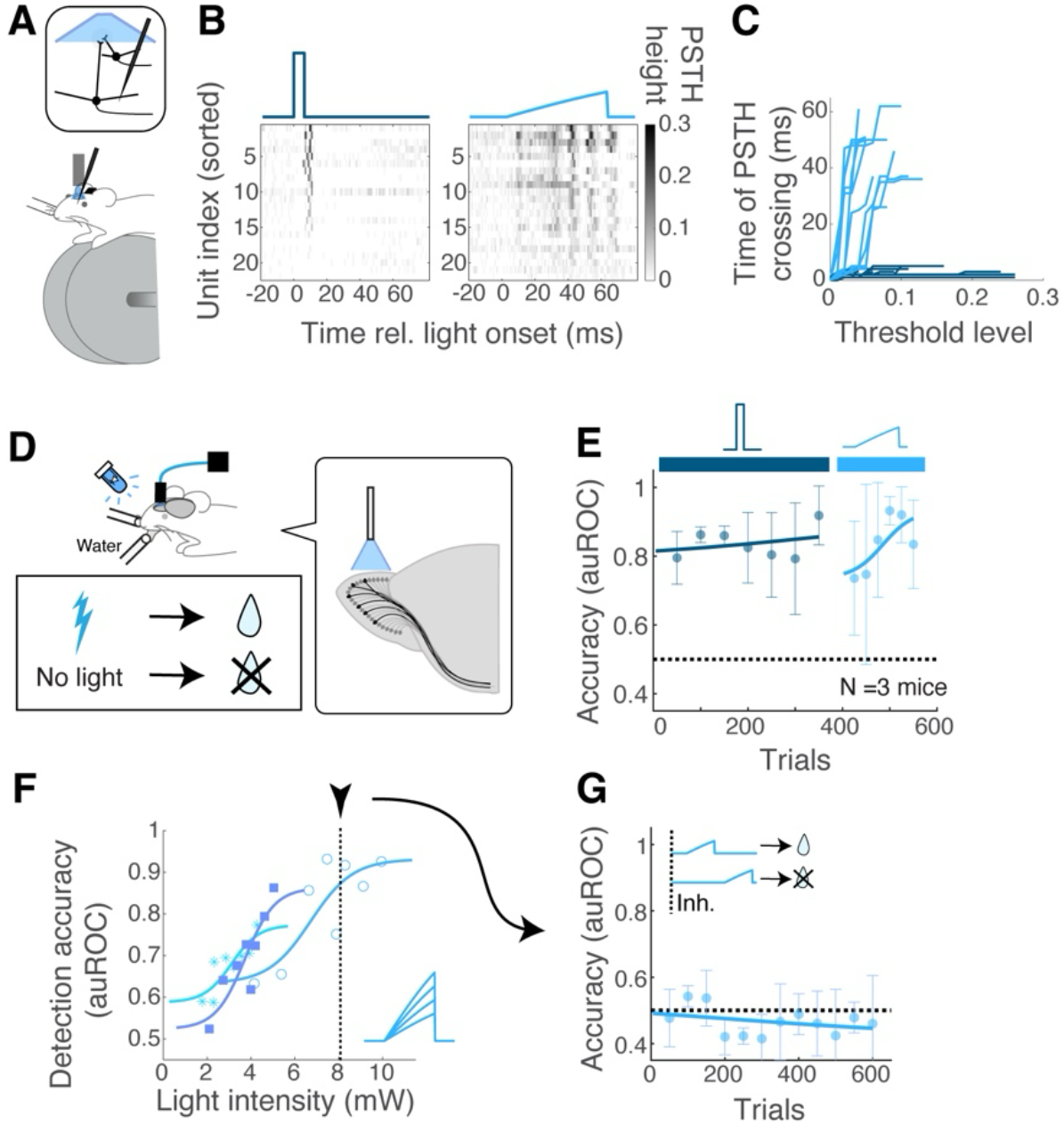
Latencies are indistinguishable with minimal OB stimulations. **A**. To achieve weaker stimulations of mitral and tufted cells, light was now presented on the olfactory bulb, through an implanted optical window. The advantage of this method is the ability to recruit neurons less synchronously. Optogenetic recruitment of mitral and tufted cells using this method was confirmed using extracellular electrophysiology in awake, head-fixed mice. **B**. Peri-stimulus time histogram of putative mitral and tufted unit activities for rectangular light waveform (dark blue, left panel) and ramp light waveform (cyan, right panel). **C**. Time when the PSTH height crossed a threshold was quantified for each unit. When units in the population are recruited synchronously and with short latency, it is characterised by an early time crossing that remains consistent for various threshold levels used. **D**. Experimental configuration for the detection task. Rewarded stimulus was presence of light presentation, and the unrewarded stimulus was its absence. **E**. Learning curve for rectangular waveform and the ramp waveform. Light intensities at the maximum were 50 mW and 12 mW for the rectangular vs. ramp waveforms. **F**. Measuring the detection threshold for the ramp waveform by changing the light intensity used. The psychometric curve between the light intensity and the detection accuracy for 3 animals (individual lines). Inset shows a schematic - ramp light at different intensities were presented within one session. **G**. Learning curve of latency discrimination for ramp light at detection threshold (8 mW). Mice were not able to distinguish the evoked timing with respect to the sniff cycle with this condition. N = 3 mice

